# Identification of a recombinant human astrovirus type 5 strain from an acute gastroenteritis outbreak in Beijing, China

**DOI:** 10.1101/2023.06.27.546739

**Authors:** Tongyao Mao, Peng Zhang, Lili Li, Mengyao Yang, Yalu Wang, Yalin Ma, Dandi Li, Jinsong Li, Zhaojun Duan

## Abstract

Two samples positive for the same strain (2103CP) of human astrovirus (HAstV) were identified during an acute gastroenteritis outbreak in Beijing, China. A full genomic analysis showed that for both samples, open reading frame 1a (ORF1a) clustered with HAstV-1, whereas ORF1b and ORF2 clustered with HAstV-5. The recombination site was detected upstream of the region where ORF1a and ORF1b overlapped. Recombinant HAstV-5 strains were reported previously in China in 2013 and 2021. Our results indicate the need for continuous surveillance of the prevalence and evolution of this recombinant HAstV-5 in China.

## Introduction

Human astrovirus (HAstV: Astroviridae) is an important pathogen of viral diarrhea in infants, young children, the elderly, and patients with immune deficiency, the first strain of which (HAstV-1) was discovered in 1975. Circulation of HAstV in human populations and the environment can lead to outbreaks. Eight serotypes, collectively named MAstV 1, are considered “classical” HAstV strains. Recently, three newly emerged HAstV types were reported: HAstV-MLB from Melbourne, Australia (MAstV 6), HAstV-VA from Virginia and other regions (MAstV 8), and HAstV-HMO of human– mink–ovine origin (MAstV 9) (Hou *et al*., 2016; Johnson *et al*., 2017). Among these strains infecting human beings, the eight classical serotypes were identified as the etiological agents in 2–9% of children hospitalized with diarrhea (Bosch *et al*., 2014).

HAstV is a nonenveloped, single-stranded RNA virus whose genome includes a 5’ noncoding region (NCR); three sequential open reading frames (ORFs) designated 1a, 1b, and 2; and an 80 nt 3’ NCR with a poly A tail at the 3’ end (Carter and Willcocks, 1996). ORF1a and ORF1b encode nonstructural proteins, and ORF2 encodes a capsid protein (Bosch *et al*., 2014). HAstV genotypes are mainly determined according to a gene sequence in the ORF2 region near the 3′ end, and the HAstV serotypes are consistent with their corresponding genotypes (Kroneman *et al*., 2013; Martella *et al*., 2014). HAstV with a single-stranded RNA genome is highly prone to mutation or recombination. Therefore, full genomic analyses of circulating HAstV strains will contribute to our understanding of the genetic diversity, evolution, and pathogenicity of the virus.

HAstV is extremely prevalent worldwide, and its genotypes vary regionally. High positivity rates of the classical HAstV serotypes have been reported in China, South America, Africa, and other regions (Wang *et al*., 2013; Xavier *et al*., 2015; Luchs *et al*., 2021). An analysis in adults in the USA showed that specific immunoglobulin G antibody positivity of classical HAstV serotypes decreased in the following order: HAstV-1 (73%), HAstV-3 (62%), HAstV-4 (52%), HAstV-5 (29%), HAstV-8 (27%), HAstV-2 (22%), HAstV-6 (8%), and HAstV-7 (8%) (Meyer *et al*., 2021). Globally, HAstV-1 is the most prevalent serotype, accounting for >50% of all recent reports (Vu *et al*., 2017). HAstV-5 is very rarely reported in China (Guo *et al*., 2010; Yu *et al*., 2020). To date, more than 90 complete genome sequences of HAstV-1 worldwide have been listed in PubMed, whereas only 14 complete genome sequences of HAstV-5 are available. Among these, 12 have been detected in developiaang countries (Vu *et al*., 2017), which is helpful for analyses of genotypic variation and health monitoring in these regions. In China, two cases of recombinant HAstV-5 were reported in 2013 and 2020, in a 3-year-old child with diarrhea and an immunocompromised patient, respectively.

This study is the first to report recombinant HAstV-5 identified from an acute gastroenteritis (AGE) outbreak that occurred in a childcare facility in Beijing, China. The genomes of two HAstV-5-positive samples (2103CP1 and 2103CP2) were obtained for molecular and evolutionary analyses. Our results provide a foundation for elucidating the full genetic variation in HAstV-5.

## Methods and Results

During an AGE outbreak, RNA was extracted from four stool samples collected from children with AGE and purified using the QIAamp Viral RNA Mini Kit (Qiagen, Hilden, Germany). A quantitative reverse-transcription polymerase chain reaction (qRT-PCR) method established by our laboratory, including an independent RT step, was performed using the SuperScript III Reverse Transcription Kit (Invitrogen, Waltham, MA, USA). This method detected HAstV along with rotavirus, norovirus, sapovirus, and intestinal adenovirus strains 40 and 41. In two samples, only HAstV was detected. The nucleic acids of these samples were amplified using the primers Mon269 and Mon270 for sequencing and preliminary genotyping (Table S1). Whole-genome cDNA was further amplified by PCR using previously published primer sets as well as those designed in this study (Table S1), with the 2 × Phanta Max Master Mix (Vazyme Biotech Co., Ltd., Nanjing, China). The PCR conditions were as follows: 3 min at 98°C, followed by 38 cycles of amplification (15 s at 98°C, 20 s of touchdown from 62°C to 54°C, and 20 s at 72°C), with a final extension for 5 min at 72°C. The 5’ and 3’ terminal sequences were amplified using the HiScript-TS 5’/3’ RACE Kit (Vazyme Biotech Co., Ltd.) according to the manufacturer’s instructions. After sequencing and assembly, the full-length sequences (6800 bp) of both samples were obtained. The genotype of each genome segment was preliminarily determined using the National Center of Biotechnology Information (NCBI) blast tool (https://blast.ncbi.nlm.nih.gov/blast.cgi), and the obtained sequences were submitted to GenBank (accession nos. OP482093 and OP482094; Table S2).

The 6800 bp genomic sequences of the HAstV-positive samples shared 100% nucleotide and 100% amino acid sequence identity, indicating that they were the same strain (2103CP). The blast results showed that this strain shared > 98.7% nucleotide and amino acid homology with recombinant HAstV-5 strains detected in China in 2013 (Fuzhou/85) and 2020 (Yu/1-CHN) (Huang *et al*. 2018; Yu *et al*. 2020). The predicted genomic structure (Fig. 1) includes three ORFs (ORF1a, ORF1b, and ORF2), as well as 85 bp NCRs at the 5’ and 3’ ends. The total length of ORF1a was 2799 bp (sites 86– 2884), encoding 932 amino acids, whereas that of ORF1b was 1548 bp (sites 2824– 4371), encoding 515 amino acids; that of ORF2 was 2352 bp (sites 4364–6715), encoding 783 amino acids. Sequence analysis showed that ORF1a and ORF1b had an overlapping region of 61 bp, whereas ORF1b and ORF2 had eight base pair repeats, and the 3’ end contained a poly A tail, highly consistent with the previously reported Chinese HAstV-5 strains Fuzhou/85 and Yu/1-CHN.

**Fig. 1.**
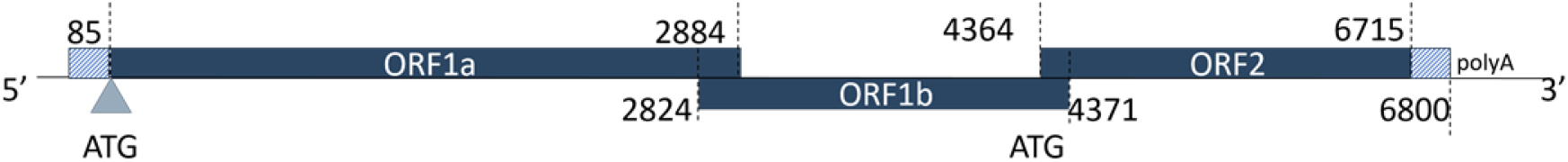
RNA genome organization of the HAstV-5 2103CP1 and 2103CP2 strains (GenBank ID: OP482093 and OP482094). The triangle represents the site of the predicted start codon (ATG).

Amino acid sequence similarity was calculated using MegAlign (DNASTAR, Madison, WI, USA). The ORF1a, ORF1b, and ORF2 regions of strain 2103CP shared 91.9–99.5%, 97.6–100%, and 96.9–99.2% amino acid identity, respectively, with the HAstV-5 reference strains. Using MEGA v7.0.26, we compared the HAstV-1/5 recombinant strains, Yu/1-CHN and Fuzhou/85, to strain 2103CP, which was found to have eight amino acid mutations in the ORF1a region (D/E15D, K34R, T61A, Q177H, A/V615A, K/E781E, A786V, and H/Y806H) and a deletion at N180. Strain 2103CP shared only 93.1% homology with the HAstV-5 strain DL030 (isolate JQ403108.1) in ORF1a but up to 98.1% amino acid identity with the HAstV-1 strain Pune/063681 (isolate JF327666.1). These results suggest that strain 2103CP may have been recombined with HAstV-1 to produce strains similar to Yu/1-CHN and Fuzhou/85.

Maximum-likelihood phylogenetic trees were constructed using MEGA v7.0.26 based on general time-reversible models (Figs. 2 and 3A, 3C) and Tamura-Nei models(Figs. 3B), with 1000 bootstrap replicates. The reference sequences were retrieved from GenBank. Strain 2103CP clustered with HAstV-5 strains including Fuzhou/85 and Yu/1-CHN from China, and with other HAstV-5 strains detected worldwide from 2013 to 2021 (Fig. 2). Strain 2103CP clustered tightly with all HAstV-5 strains, including DL030 strains, in ORF1b and ORF2, whereas in ORF1a, strain 2103CP clustered tightly with the HAstV-1 strain Pune/063681. This result further suggests that strain 2103CP may be a recombinant strain of HAstV-1/5, similar to strains Fuzhou/85 and Yu/1-CHN.

**Fig. 2.**
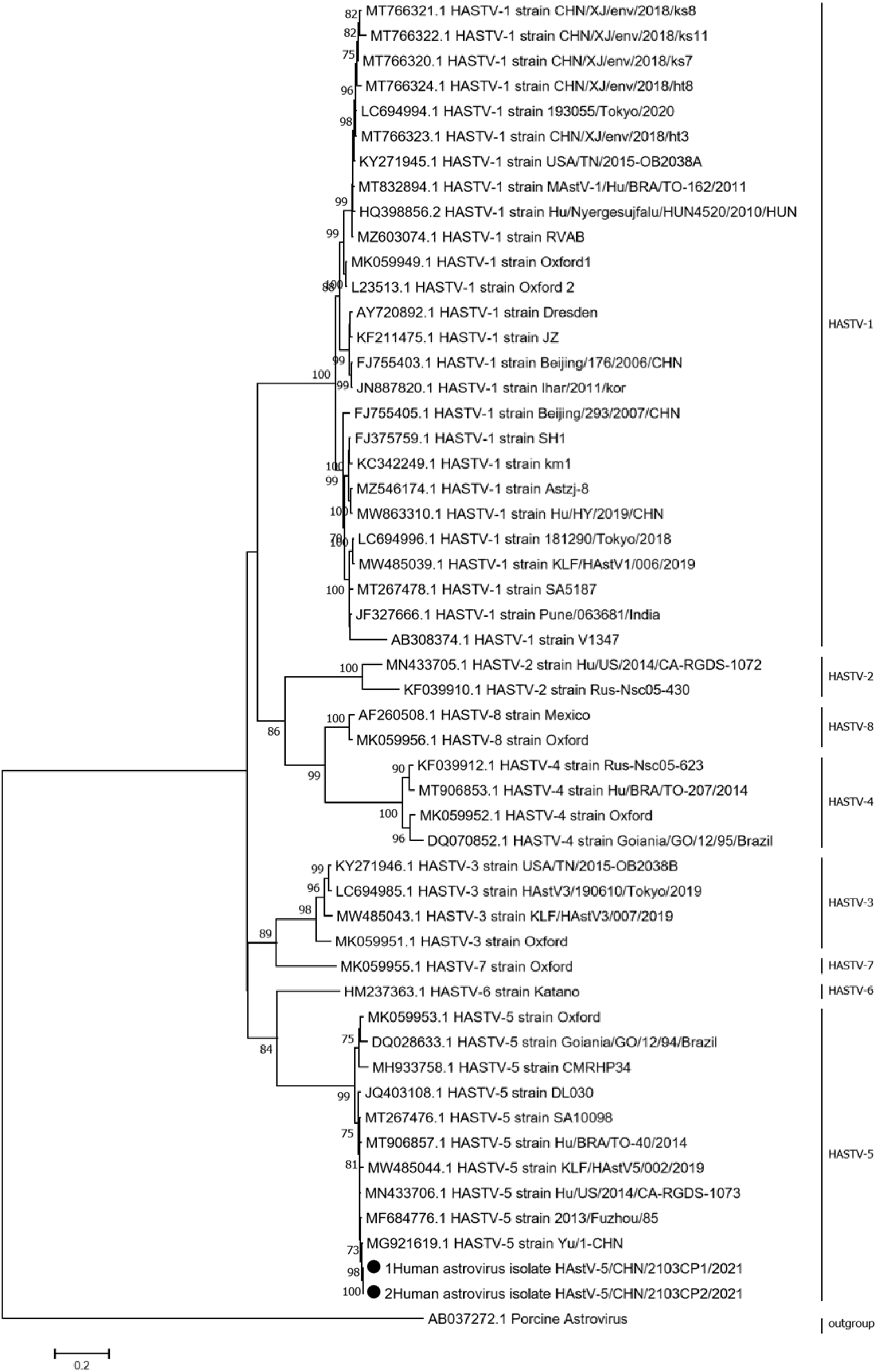
Phylogenetic analysis of the entire genome of circulating human astrovirus 1-8, which is seroprevalent worldwide. A GTR+G nucleotide substitution model was used to construct the phylogenetic tree. The porcine AB037272 strain was used as the outgroup. Dots indicate strains 2013CP1 and 2013CP. Bootstrap values (1000 replicates) ≥ 70% are shown.

**Fig. 3.**
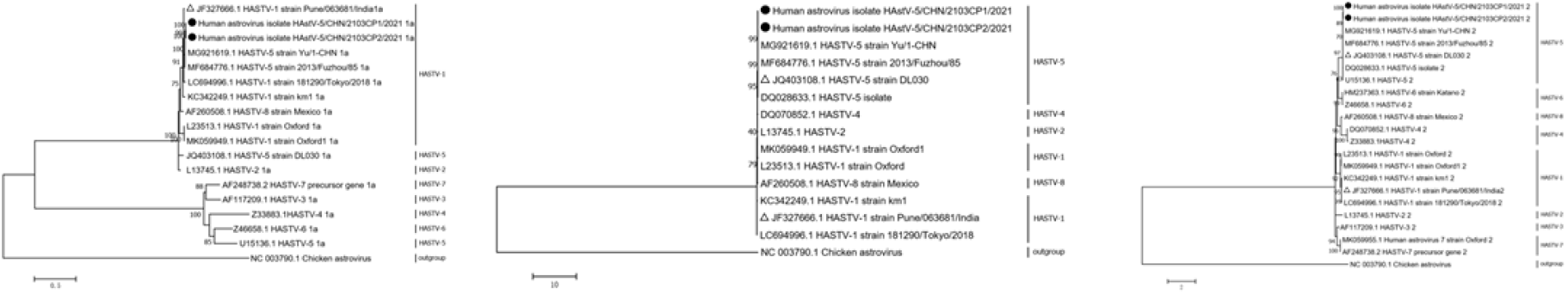
Phylogenetic trees based on the nucleotide sequences of the ORF1a (A), ORF1b (B), and ORF2 (C) regions of strains 2013CP1 and 2013CP2 and other representative astrovirus strains. The maximum-likelihood tree was constructed based on the Tamura-Nei (B) and General Time Reversible models (A, C) models and with 1000 bootstrap replicates. Dots indicate strains 2013CP1 and 2013CP; triangles indicate the strain used for the recombination analysis. Bootstrap values (1000 replicates) ≥ 70% are shown.

SimPlot analysis showed a putative crossover at 2687 nt in strain 2103CP (Fig. 4). Nucleotide sequences before and after the crossover point were highly similar to those of HAstV-1 strain Pune/063681 and HAstV-5 strain DL030. The recombination site was detected upstream of the region where ORF1a and ORF1b overlapped, as reported for strains Fuzhou/85 and Yu/1-CHN, indicating that these novel HAstV-5 recombinants share a common ancestor, through continuous transmission in various regions of China.

**Fig. 4.**
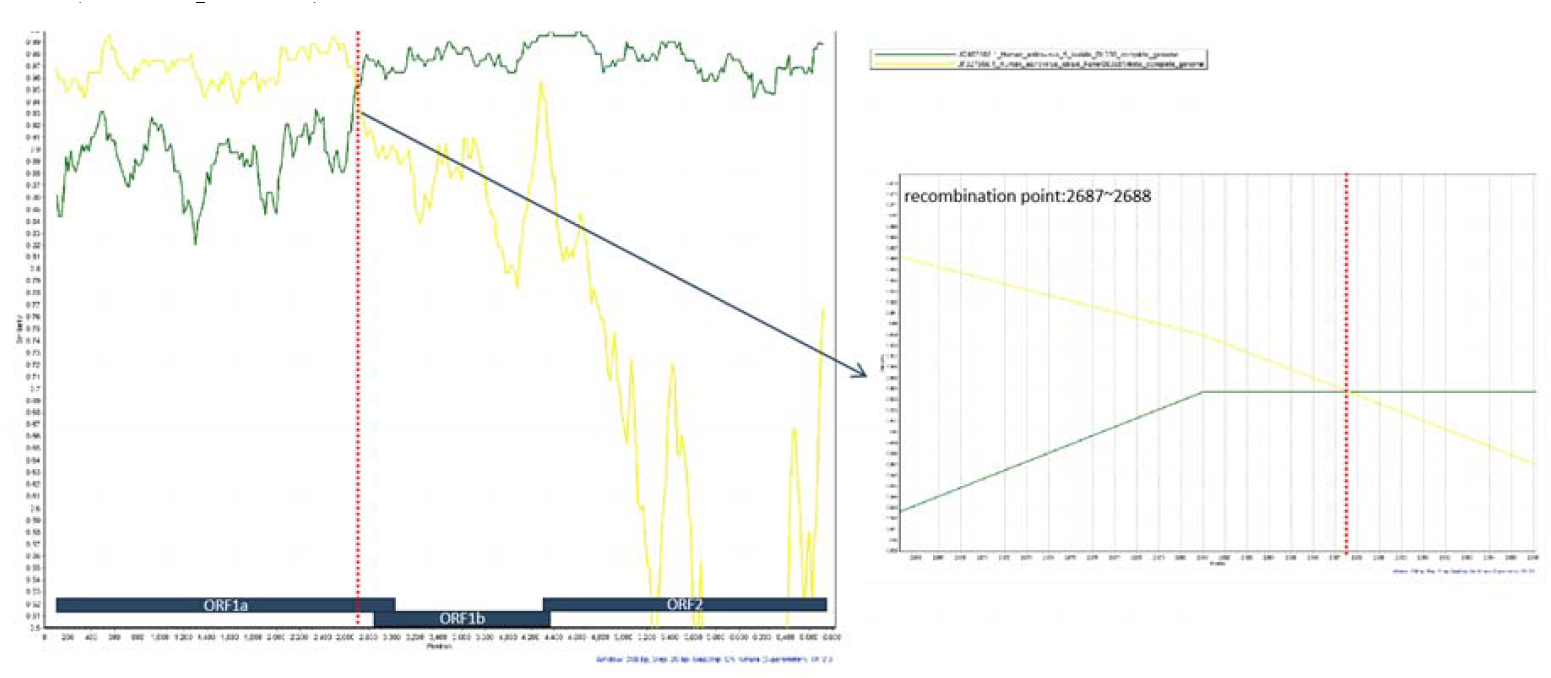
Whole genome recombination analysis of human stellate viruses 2103CP1 and 2103CP2 using Simplot software. The yellow and green curves represent the identities of 2103CP1 compared to HASTV-1 strain Pune/063681/India and strain HASTV-5 DL030, respectively.

The breakpoint of the novel HAstV-5 recombinants occurred upstream of the ORF1a/ORF1b junction region. Previous studies have predominantly reported the ORF1b/ORF2 junction as a recombination hotspot (Walter *et al*., 2001; Medici *et al*., 2015; Ha *et al*., 2017; Wei *et al*., 2023). The ORF1b–ORF2 overlap region contains sequences that are conserved in HAstV (Méndez and Arias, 2013; Medici *et al*., 2015), which may increase the probability of homologous recombination, allowing structural protein exchange (Wohlgemuth *et al*., 2019). Recombination is common in many RNA virus families (Worobey and Holmes, 1999), increasing their viral diversity and survival rates. Recent studies have demonstrated that recombination is an intrinsic defense in RNA viruses against host antiviral interference (Aguado *et al*., 2018). Thus, genome sequencing is an important and necessary tool for monitoring HAstV evolution.

## Discussion

With the rapid development of virus detection technology and public health awareness, HAstV has become recognized as an important agent of diarrhea that is associated with food- and water-borne pollution. Our findings demonstrate the circulation and continuous prevalence of recombinant HAstV-5 strains with novel ORF recombination sites in China. Limitations of this study include that environmental samples, including food and water, were not collected and the source of the AGE outbreak was not explored. Further epidemiological research is necessary to monitor the novel recombinant strain within the population, particularly during AGE outbreaks, to elucidate the transmission and pathogenesis of HAstV, as well as its disease burden and molecular characteristics. The resulting genomic data are anticipated to improve diagnosis techniques and vaccine development of HAstV.

## Data availability statement

The datasets generated and/or analyzed during the current study are available from the corresponding author on reasonable request.

## Author contributions

Tongyao Mao, Peng Zhang, Jinsong Li and Zhaojun Duan were involved in the design of this study. Mengyao Yang, Yalu Wang, Yalin Ma performed RNA extraction, sequencing and sequences classification. Tongyao Mao and Lili Li performed molecular and phylogenetic analyses. Dandi Li participated in sample collection. Tongyao Mao wrote the manuscript. Zhaojun Duan was responsible for the critical revision of the manuscript. All authors have received the final version of the manuscript.

## Ethics Exemption Declarations

This study uses strains obtained from the remaining abandoned samples after the in-hospital test of children with diarrhea. The ethics committee of the National Institute of Viral Disease Control and Prevention did not require the study to be reviewed or approved by an ethics committee because this project is the reuse of these samples, which will not cause any harm to the subjects. The personal information involved in the research records and results are identified by numbers, so it will not cause the disclosure of personal privacy.

## Appendix A. Supplementary data

Supplementary material related to this article can be found, in the online version, at doi:https://doi.org/xxx/xxx.

## Supplementary material

**Table S1.**
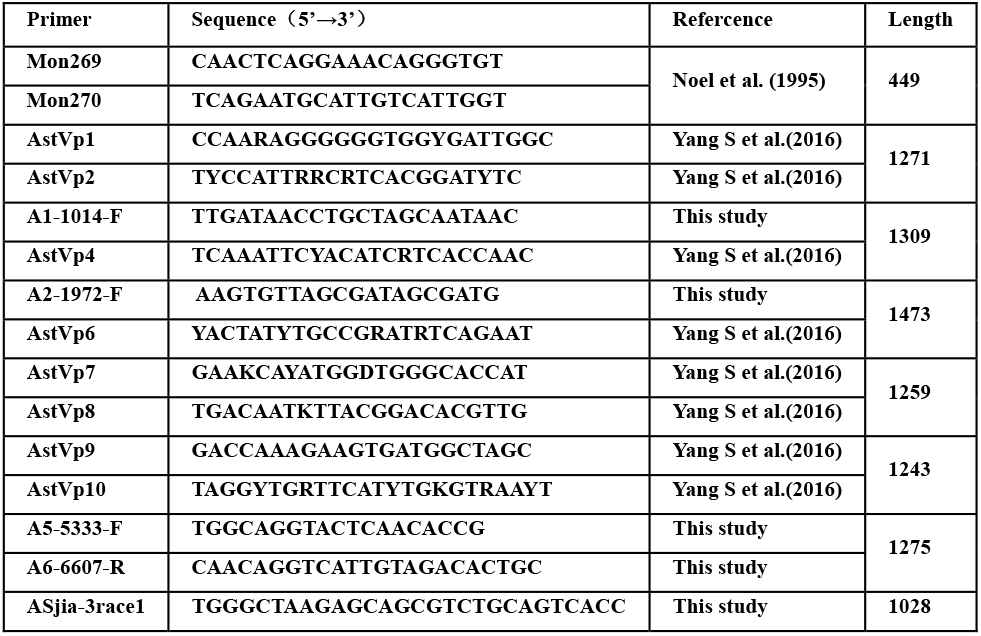
Primers used in this study

**Table S2.** Information about the two HAstV-5 strains examined in this study.

| Strain | Collection date | Location | Genebank |
| --- | --- | --- | --- |
| 2103CP1 | 2021 | Beijing | OP482093 |
| 2103CP2 | 2021 | Beijing | OP482094 |

## References

Aguado LC, Jordan TX, Hsieh E, Blanco-Melo D, Heard J, Panis M, Vignuzzi M, tenOever BR. 2018. Homologous recombination is an intrinsic defense against antiviral RNA interference. Proceedings of the National Academy of Science 115:E9211–E9219.

Bosch A, Pintó RM, Guix S. 2014. Human astroviruses. Clinical Microbiology Reviews 27:1048–1074.

Carter MJ, Willcocks MM. 1996. The molecular biology of astroviruses. Archives of Virology. Supplementum 12:277–285. DOI: 10.1007/978-3-7091-6553-9_30

Guo L, Xu X, Song J, Wang W, Wang J, Hung T. 2010. Molecular characterization of astrovirus infection in children with diarrhea in Beijing, 2005–2007. Journal of Medical Virology 82:415–423. DOI: 10.1002/jmv.21729

Ha H-J, Lee SG, Cho H-G, Jin J-Y, Lee JW, Paik S-Y. 2016. Complete genome sequencing of a recombinant strain between human astrovirus antigen types 2 and 8 isolated from South Korea. Infection, Genetics and Evolution 39:127–131. DOI: 10.1016/j.meegid.2016.01.017

Hou M, Liang X, Tan L, Liu P, Zhao W. 2016. Outbreak of human astrovirus 1 lineage 1d in a childcare center in China. Virologica Sinica 31:258–261.

Huang Z, Wu B, Chen F, Huang Y, Weng Y. 2018. Whole-genomic analysis of a recombinant HAstV-5 astrovirus identified in Fujian, China. Chinese Journal of Virology 34:16–21. DOI: 10.13242/j.cnki.bingduxuebao.003288

Johnson C, Hargest V, Cortez V, Meliopoulos VA, Schultz-Cherry S. 2017. Astrovirus pathogenesis. Viruses 9:22.

Kroneman A, Vega E, Vennema H, Vinje J, White PA, Hansman G, Green K, Martella V, Katayama K, Koopmans M. 2013. Proposal for a unified norovirus nomenclature and genotyping. Archives of Virology 158:2059–2068. DOI: 10.1007/s00705-013-1708-5.

Luchs A, Tardy K, Tahmasebi R, Guadagnucci Morillo S, de Pádua Milagres FA, Dos Santos Morais V, Brustulin R, da Aparecida Rodrigues Teles M, de Azevedo LS, de Souza EV et al. 2021. Human astrovirus types 1, 4 and 5 circulating among children with acute gastroenteritis in a rural Brazilian state, 2010–2016. Archives of Virology 166:3165–3172. DOI: 10.1007/s00705-021-05206-8

Martella V, Pinto P, Tummolo F, De Grazia S, Giammanco GM, Medici MC, Ganesh B, L’Homme Y, Farkas T, Jakab F, Bányai K. 2014. Analysis of the ORF2 of human astroviruses reveals lineage diversification, recombination and rearrangement and provides the basis for a novel sub-classification system. Archives of Virology 159:3185–3196. DOI: 10.1007/s00705-014-2153-9

Medici MC, Tummolo F, Martella V, Banyai K, Bonerba E, Chezzi C, Arcangeletti MC, De Conto F, Calderaro A. 2015. Genetic heterogeneity and recombination in type-3 human astroviruses. Infection, Genetics and Evolution 32:156–160. DOI: 10.1016/j.meegid.2015.03.011

Méndez E, Arias C. 2013. Astroviruses. In: Knipe DM, Howley PM, eds. Fields Virology Volume 1, 6th ed. Philadelphia, PA: Lippincott Williams & Wilkins. pp. 609–628.

Meyer L, Delgado-Cunningham K, Lorig-Roach N, Ford J, DuBois RM. 2021. Human astrovirus 1–8 seroprevalence evaluation in a United States adult population. Viruses 13:979. DOI: 10.3390/v13060979

Vu D-L, Bosch A, Pintó RM, Guix S. 2017. Epidemiology of classic and novel human astrovirus: Gastroenteritis and beyond. Viruses 18:33. DOI: 10.3390/v9020033.

Walter JE, Briggs J, Guerrero ML, Matson DO, Pickering LK, Ruiz-Palacios G, Berke T, Mitchell DK. 2001. Molecular characterization of a novel recombinant strain of human astrovirus associated with gastroenteritis in children. Archives of Virology 146:2357–2367.

Wang Y, Li Y, Jin Y, Li DD, Li X, Duan ZJ. 2013. Recently identified novel human astroviruses in children with diarrhea. Emerging Infectious Diseases 19:1333–1335. DOI: 10.3201/eid1908.121863.

Wei H, Kumthip K, Khamrin P, Yodmeeklin A, Jampanil N, Phengma P, Xie Z, Ukarapol N, Ushijima H, Maneekarn N. 2023. Triple intergenotype recombination of human astrovirus 5, human astrovirus 8, and human astrovirus 1 in the open reading frame 1a, open reading frame 1b, and open reading frame 2 regions of the human astrovirus genome. Microbiology Spectrum e0488822. DOI: 10.1128/spectrum.04888-22

Wohlgemuth N, Honce R, Schultz-Cherry S. 2019. Astrovirus evolution and emergence. Infection, Genetics and Evolution. 69:30–37. DOI: 10.1016/j.meegid.2019.01.009.

Worobey M, Holmes EC. 1999. Evolutionary aspects of recombination in RNA viruses. Journal of General Virology 80:2535–2543.

Xavier MdPTP, Carvalho Costa FA, Rocha MS, de Andrade JdSR, Diniz FKB, de Andrade TR, Miagostovich MP, Leite JPG, de Mello Volotão E. 2015. Surveillance of human astrovirus infection in Brazil: The first report of MLB1 astrovirus. PLoS ONE. 10:e0135687. DOI: 10.1371/journal.pone.0135687.

Yu J-M, Wang Z-H, Liu N, Zhang Q, Dong Y-J, Duan Z-J. 2020. Complete genome of a novel recombinant human astrovirus and its quasispecies in patients following hematopoietic stem cell transplantation. Virus Research 288:198138. DOI: 10.1016/j.virusres.2020.198138

